# Evolution of resistance in vitro reveals a novel mechanism of artemisinin activity in *Toxoplasma gondii*

**DOI:** 10.1101/746065

**Authors:** Alex Rozenberg, Madeline R. Luth, Elizabeth A. Winzeler, Michael Behnke, L. David Sibley

## Abstract

Artemisinins are effective against a variety of parasites and provide the first line of treatment for malaria. Laboratory studies have identified several mechanisms for artemisinin resistance in *Plasmodium falciparum*, including mutations in Kelch13 that are associated with delayed clearance in some clinical isolates, although other mechanisms are likely involved. To explore other potential mechanisms of resistance in parasites, we took advantage of the genetic tractability of *T. gondii*, a related apicomplexan parasite that shows moderate sensitivity to artemisinin. Resistant populations of *T. gondii* were selected by culture in increasing drug concentrations and whole genome sequencing identified several non-conservative point mutations that emerged in the population and were fixed over time. Genome editing using CRISPR/Cas9 was used to introduce point mutations conferring amino acids changes in a serine protease homologous to DegP and a serine/threonine protein kinase of unknown function. Single and double mutations conferred a competitive advantage over wild type parasites in the presence of drug, despite not changing EC_50_ values. Additionally, the evolved resistant lines showed dramatic amplification of the mitochondrial genome, including genes encoding cytochrome b and cytochrome oxidase I. Consistent with prior studies in yeast and mammalian tumor cells that implicate the mitochondrion as a target of artemisinins, treatment of wild type parasites with artemisinin decreased mitochondrial membrane potential, and resistant parasites showed altered morphology and decreased membrane potential. These findings extend the repertoire of mutations associated with artemisinin resistance and suggest that the mitochondrion may be an important target of inhibition in *T. gondii*.

**Significance:** Artemisinins provide important therapeutic agents for treatment of malaria and have potential for use in other infections and in cancer. Their use is threatened by the potential for resistance development, so understanding their mechanism of action and identifying genetic changes that alter sensitivity are important for improving clinical outcomes. Our findings suggest that mutations in novel targets can contribute to the emergence of parasites with increased tolerance to artemisinin treatment and that such mutations can confer a fitness advantage even in the absence of a notable shift in EC_50_. Our findings also support the idea that inhibition of mitochondrial function may be an important target in *T. gondii*, as previously suggested by studies in yeast and human cancer cells.

## Introduction

*Toxoplasma gondii* is a widespread parasite of animals that also frequently causes zoonotic infections in people (1). Serological testing suggests that ∼1/3 of the global human population is chronically infected (2), although such infections are typically well controlled by the immune response following mild clinical symptoms in the acute phase (3). Chronic infections are characterized by semi-dormant tissue cysts containing bradyzoites, which divide asynchronously and sporadically (4). Current therapies and induction of a potent immune response are insufficient to clear these stages and instead they undergo slow turnover to sustain infection over time. Re-emergence of chronic infection in patients that are immunocompromised results in serious disease (5, 6), especially when this occurs in the central nervous system where tissue cysts most often infect neurons (7). Toxoplasmosis is also a threat due to risk of congenital infection, especially in resource-limited regions (8). More severe disease has been reported in some regions of South America where infections are associated with ocular disease in otherwise healthy individuals (9). Toxoplasmosis is typically treated by anti-folates using a combination of pyrimethamine-sulfadiazine or trimethoprim-sulfamethoxazole, although pyrimethamine has also been used in combination with clindamycin, azithromycin, and atovaquone (also used as monotherapy) (10). With the exception of congenital infection, infections are not commonly transferred from human-to-human, so emergence of drug resistance is rarely a problem. However, several examples of clinical failure have been reported in patients given atovaquone as a monotherapy (11, 12).

Combination therapies including artemisinin (ART) are the first line of treatment against malaria (13). Since their original discovery (14), a wide variety of analogs have been made to improve solubility and other drug-like properties, including artemether, artusunate and dihydroartemisinin, which is the active form *in vivo* (15). ART therapies are fast acting and effective across multiple life cycle stages; however, they have short half lives *in vivo* and must be coupled with longer lasting partner drugs to reduce the risk of resistance development (13). ART derivatives share an endoperoxide bond that is essential for activity, demonstrated by the fact that the derivative deoxyartemisinin loses antimalarial activity (16). More recently, completely synthetic analogs containing a similar ozonide group have been developed and some of these have increased half lives *in vivo* (17). ART analogs are also effective in blocking growth of *T. gondii in vitro* (18-20) and controlling infection *in vivo* in murine models of toxoplasmosis (21, 22). Although analogs show similar potency ranking, they are effective against *T. gondii* at ∼50-fold higher concentrations when compared to *P. falciparum* (20, 23). This major difference in sensitivity may be due to the fact that *P. falciparum* digests hemoglobin in the food vacuole and releases heme in high concentrations (24). The majority of heme is detoxified into a crystalline form called hemozoin, and yet it is likely that elevated reduced iron (Fe^2+^) is present in the food vacuole. One potential mechanism is that free Fe^2+^ activates the endoperoxide bond (25), resulting in formation of adducts to a variety of protein and lipid targets that result in parasite death (26, 27). There is no analogous process to hemoglobin digestion in *T. gondii*, which may explain its lower sensitivity to ART. Instead, it is possible that heme derived from sources such as mitochondria plays an important role in activating ART in *T. gondii*, similar to yeast and tumor cell models, as discussed below.

Treatment failure in *P. falciparum* patients treated with ART was first reported in western Cambodia as a delayed clearance phenotype (28). However, such isolates are not “resistant” by the classical definition of shifted EC_50_; rather they show delayed clearance times compared to the normal rapid drop in parasitemia following ART treatment (13). The delayed clearance phenotype from many SE Asian isolates correlates with outgrowth after high dose treatment of ring stages *in vitro*, the basis for the Ring stage Survival Assay (RSA) (29). Selection for ART-resistant *P. falciparum* lines in the laboratory by step-wise increases in drug concentration led to identification of mutations in a Kelch13 (K13) ortholog (30). Subsequent reverse genetic experiments confirmed that these mutations were capable of conferring resistance in a wild type background (31). Additionally, a number of K13 mutations were found in high abundance in SE Asia where they correlated with RSA sensitivity (30). Several mechanisms have been proposed to explain the role of K13 in ART resistance, including an enhanced cell stress response leading to upregulation of the unfolded protein response, decreased ubiquitination, and increased cell survival (32). Consistent with this model, inhibitors of the proteasome synergize with artemisinin (33, 34). However, other studies have reported the development of delayed clearance or dormancy phenotypes in *P. falciparum*-infected patients without associated K13 mutations (35, 36). Furthermore, genetic crosses between parasite lines bearing sensitive vs. resistant K13 mutations demonstrated a correlation with the RSA assay *in vitro* but not with clearance of blood infection following treatment of infected *Aotus* monkeys, suggesting other mechanisms are responsible *in vivo* (37). Separately it has been proposed that ART inhibits PI_3_K, and that K13 mutations that confer resistance led to increased PI_3_K activity and increased IP_3_ and membrane vesicles that may confer protection to ring stages (38). In a separate study involving *in vitro* selections, resistance-conferring mutations were identified in a number of genes including the actin-binding protein coronin(39). In summary, there are a number of distinct mechanisms that can lead to decreased sensitivity of *P. falciparum* to ART, which complicates efforts to track the development of resistance in the field.

A number of specialized artemisinin derivatives have been synthesized and promoted as potential treatments for cancer as they are active against a wide variety of tumor cells *in vitro* (40). The activities of the common ART derivatives against tumor cell lines are in the low micromolar range, which is comparable to their activity in *T. gondii* but much higher than what is typically observed against *P. falciparum*. The ART mechanism of action against tumor cells is also uncertain, but has been linked to oxidative stress, DNA damage, decreased cell replication, and activation of cell death pathways including apoptosis (41, 42). Studies in Baker’s yeast (*S. cerevisiae*) implicate the mitochondria as a target of ART, and consistent with this model, growth on fermentable carbon sources results in insensitivity as the mitochondria is dispensable under these conditions (43). Treatment of yeast with ART results in mitochondrial membrane depolarization, suggesting that ART is activated by the electron transport system which leads to oxidative stress and membrane damage (44). Inhibition of mitochondrial function in tumor cells treated with ART derivatives has also been shown to promote cell death due to apoptosis (45). In human tumor cells, it is proposed that mitochondrial heme activates the endoperoxide group of artemisinin and interaction with the electron transport chain generates reactive oxygen species leading to cell death (46).

Here we used the versatility of *T. gondii* as a genetic system to explore potential mechanisms of resistance to ART. *Toxoplasma* can be propagated asexually for long periods of time and single cell clones can be readily isolated, cryopreserved, and expanded. Multiple high-quality genome assemblies of the haploid ∼ 65 Mb genome are available from a diverse set of strains (47) and efficient systems are available for reverse genetics using CRIPSR/Cas9 (48). Previous studies have shown that ART resistance can be selected in chemically-mutagenized *T. gondii* parasites (49), although the molecular mechanism for the modest shift in EC_50_ observed in these resistant clones was not defined. Chemical mutation introduces numerous changes in the genome, confounding analysis of those that might confer resistance. Consequently, we used a natural evolution process to select for mutations that would confer increased survival in ART, while monitoring the background mutation rate in the absence of selection in parallel. We applied selective pressure by growing parasites for many generations in increasing concentrations of ART and were able to obtain resistant populations as evidenced by increased EC_50_ values. Whole genome sequencing identified several single nucleotide polymorphisms that arose in the resistant populations and reached fixation over time. Reverse genetic approaches based on CRISPR/Cas9 gene editing were used to show that these mutations confer increased competitive advantage over wildtype, consistent with enhanced tolerance, despite the fact that they did not alter EC_50_ values. Additionally, the resistant populations showed stable changes in the mitochondrion, which may contribute to resistance. These findings define novel molecular changes associated with ART resistance in *T. gondii*, which may be informative for understanding the mechanism of action of this important class of compounds.

## Results

### Estimating the background mutation rate in *T. gondii*

Prior to selecting for resistant parasites, we wanted to establish the background mutation rate of *T. gondii* during a continuous *in vitro* passage under standard laboratory conditions. Because lines that have been cultured for years in the laboratory are likely a mixture of genotypes that contain both private and shared single nucleotide polymorphisms (SNPs), as well as potential copy number variations (CNVs), we first cloned the starting population by limiting dilution on monolayers of human foreskin fibroblasts (HFF cells). We obtained clones from the common laboratory RH strain, referred to herein as the 5D and 12B lines, and grew them by sequential passage in HFF cells for 365 days, sub-culturing them every 2-3 days (Fig. 1A). During *in vitro* passage in this format, the total population parasite size of a given culture expands to ∼ 2×10^7^ over a time frame of 2-3 days. The population is then reduced by 1:10 or 1:20 when a fraction of the growing culture is used to inoculate a new flask of previously uninfected HFF cells. At the time points of 182 and 365 days, separate subclones from the growing populations were isolated by limiting dilution, and these were named with the prefix of the parental line followed by a clone designation (Fig. 1A). Genomic DNA extracted from these clones and their respective parents were whole genome sequenced using Illumina technology. The short reads were mapped with high stringency to the reference genome of the ME49 strain of *T. gondii* (www.ToxoDB.org) using a specially developed pipeline (see methods). We then identified and compared SNP frequencies between the reference parental clones (i.e. 12B and 5D) and the subclones that were obtained from these lines following passage. Overall the number of mutations detected in the clones were quite low with a range from 0 – 8 SNPs observed per clone (*Datasets* S1, S2).

**Fig 1.**
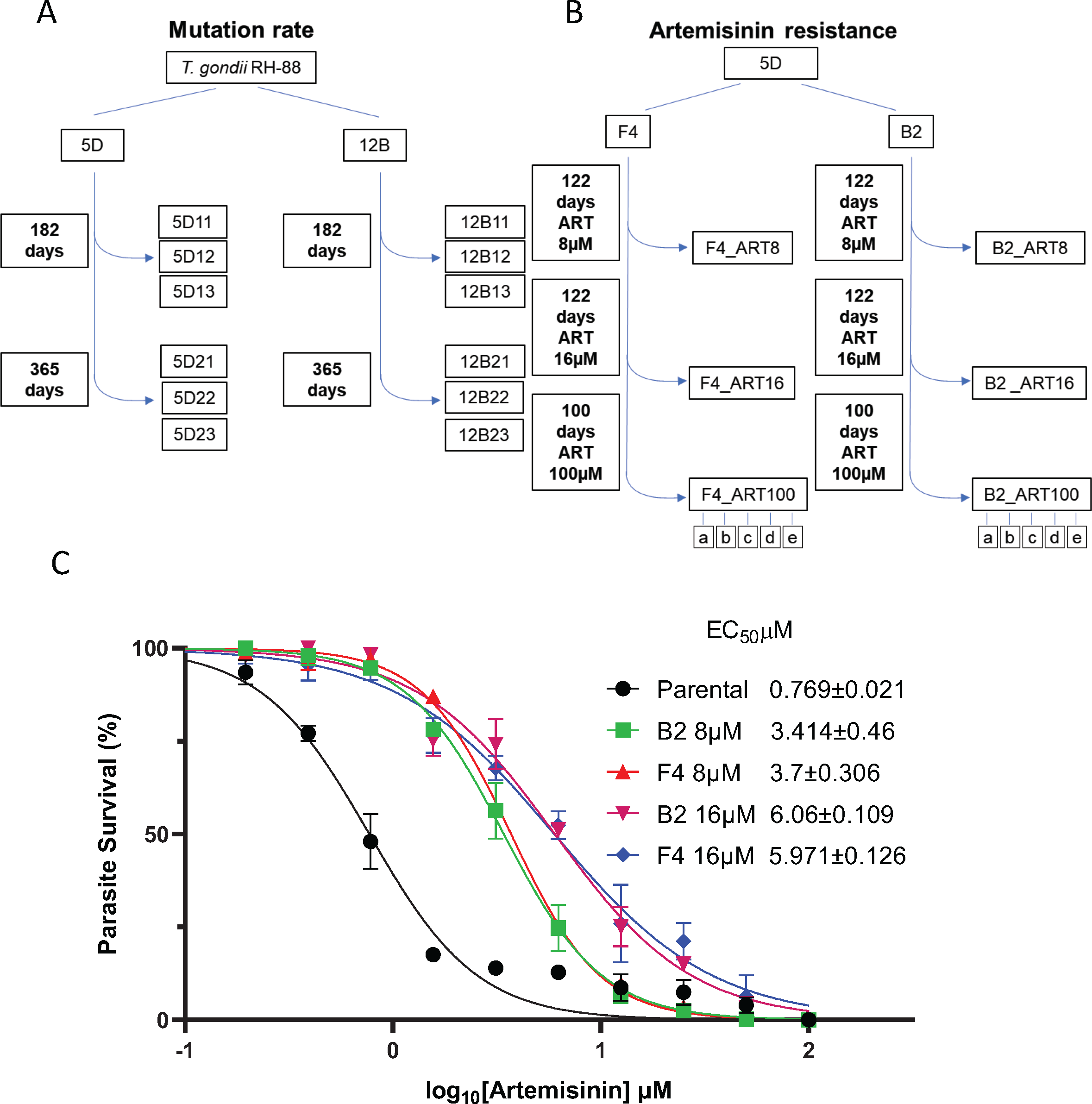
Generation of artemisinin (ART) resistant parasite populations. A) Schematic for growth of parasites used to determine the background mutation rate. Two initial RH88 strain clones (5D and 12B) were sequentially passed in T25^2^ flasks containing HFF monolayers every 48 h. The clones were subcloned by limiting dilution at 182- and 365-day time points. The genomes of the parental strain clones (5D and 12B) and the various subclones were sequenced using Illumina technology. B) Schematic for selection of ART resistant parasites. The parasite clone (5D) was further subcloned into two lines (F4 and B2) and subjected to 8 μM ART during sequential *in vitro* passages. The ART concentration was increased to 16 μM and 100 μM at the intervals shown. Populations were frozen for further analysis at the 8 μM and 16 μM levels and clones (a-e) were made after passage in 100 μM. C) Dose response curves for inhibition of *T. gondii* parental and ART adapted B2 and F4 lines growth in response to increasing concentration of ART. Parasite growth was monitored by measurement of lactate dehydrogenase (LDH) release from host cells as a consequence of rupture. Data presented as percent LDH release (% LDH) normalized to growth in naïve (untreated) HFF cells. Shown are three biological replicates each with technical replicates (n=9) ± SE. EC_50_ values were determined using non-linear regression analysis as a sigmoidal dose–response curve with variable slope. The EC_50_ data are presented as the average of three biological replicates (i.e., separate EC_50_ titrations) each with three technical replicates.

SNP frequency was used to derive approximate rates of mutation based on the assumption that the parasite doubles every ∼6 hr (four generations per day) (50) and that the composite genome is roughly 6.5 x 10^6^ bp (47). We calculated the rates for mutation in individual clones with the assumption that the mutants arose spontaneously and that the rates of mutation and growth were constant during the course of the experiment (*Datasets* S1, S2). Based on these calculations, the average rates over the course of the 365 day culture period were 6.62 x 10^-11^ mutations / bp / generation for the 5D clones and 4.71 x 10^-11^ mutations / bp / generation for the 12B clones. We also estimated the composite mutation rate for both sets of clones by treating the frequency of SNPs as mutants in a classic fluctuation analysis yielding a combined rate of 5.79 x 10^-11^ (*Datasets* S1, S2). Since both methods give highly similar rates, we used the later value rounded to 5.8 x 10^-11^ for subsequent analysis.

When comparing the types of mutations found in each of the lines, we observed approximately equal number of variants occurring in coding and noncoding regions (19 coding vs. 13 noncoding in 5D and 10 coding vs. 13 noncoding in 12B). However, within coding regions there was a marked increase in nonsynonymous mutations vs. synonymous mutations with a dN/dS ratio of 2.8 for 5D-derived clones and dN/dS ratio of 9 for 12B-derived clones. These high dN/dS ratios suggest strong selective pressure during the culture, although these clones were grown under standard laboratory conditions. In addition, there were some mutations that had apparently reached fixation as they were common to multiple clones from a particular line. For example, in clones from the 12B line, there was a missense mutation in a hypothetical protein that was found in all 6 clones (*Dataset* S1). Similarly, in clones from line 5D, there were two frameshift mutations in hypothetical proteins, one missense mutation in a hypothetical protein, and one intronic mutation that occurred in multiple clones (*Dataset* S2).

### Establishment of ART-resistant mutants of *T. gondii*

Based on the background mutation rate, we designed an experimental protocol using sequential *in vitro* culture of relatively large parasite populations in combination with stepwise increases in drug concentrations to generate parasites with elevated ART resistance. To initiate this trial, we recloned the 5D line to derive two parental lines designated as B2 and F4 and cultured them in HFF cells supplemented with 8 µM ART as a starting point for selecting parasite populations (Fig. 1B). Although growth was initially delayed, as evident from the time required to lyse the host cell monolayer, after 122 days of sequential passage at 8 µM ART, both lineages exhibited reproducible growth and rapidly lead to lysis of the monolayer within 2-3 days of inoculation. At this time point, we cryopreserved each line for future study; these lines are referred to as F4_ART8 and B2_ART8 (Fig. 1B). We then increased the ART concentration by two-fold to 16 µM and continued to passage both lines with drug until attaining the phenotype of reproducible growth and infectivity, which took approximately 122 days. At this time point, we cryopreserved each line for future study; these lines are referred to as F4_ART16 and B2_ART816 (Fig. 1B). We then increased the concentration of ART to 100 µM and passaged the lines for an additional 100 days, after which they showed normal growth. At this stage, we generated subclones from each of the F4 and B2 lines gown at 100 µM ART by limiting dilution (Fig. 1B).

### Identification of candidate genes associated with ART resistance

To determine the extent of resistance that had developed during selection, we tested the 8 µM and 16 µM populations from both the B2 and F4 lines across a dilution series of ART and calculated their EC_50_ for growth inhibition. The evolved populations showed a ∼3 and ∼6 fold increase in EC_50_ in the 8 µM and 16 µM resistant populations, respectively (Fig. 1C). To analyze genetic changes that might have conferred enhanced ART resistance, genomic DNA from all three of the selected populations from both clones (i.e. 8, 16 and 100 µM) was extracted and subjected to whole genome sequencing. Mapping of the reads to the reference genome identified nonsynonymous mutations in the selected populations that were not present in the parental lines. The background rate of mutation is 2-3 fold higher than in non-selected lines and a number of these are found in a majority of clones, albeit typically not in coding regions (Datasets S3, S4). Analysis of the mutation frequency in these clones, including those mutations in DegP or Ark1, gave a rate of 1.39 x 10^-10^ for the B2 clones (Dataset S3) and 9.77 x 10^-11^ for the F4 clones (Dataset S4). Several of the changes were only detected in one of the two selected lines. For example, mutations were identified in two candidate genes in selected lines derived only from the B2 line and in five candidate genes derived only from the F4 line (*SI Appendix*, Table S1). The majority of these changes were also only found at the highest level of selection (i.e. 100 µM ART), where they were present at allele frequencies of > 90% (*SI Appendix*, Table S1). Because they were not common to both lines, we did not pursue them further, although it remains possible that they contributed to the observed ART resistance phenotypes.

In addition, we detected nonsynonymous changes in two genes that were common to the ART resistant populations derived from both lines. The first of these was in the gene TGME49_290840, which is annotated in ToxoDB as a serine protease (Table 1). It has high similarity to DegP in *P. malariae* (expect value based on BLASTP = 2e-103; XP_028860031.1) and is referred to here as a DegP orthologue. This DegP ortholog is different from a previously named DegP-like serine protease that is localized in the rhoptry of *T. gondii* (51). We consider TGME49_290840 to be the authentic mitochondrial DegP ortholog based on the presence of a degP-htrA superfamily motif and detection in the mitochondrial proteome (52). The *T. gondii* DegP orthologue was mutated in a different position in each of the F4 and B2 lines, although both non-conservative mutations resided in the PDZ2 domain (DegP-F4-E821Q and DegP-B2-G806E) (Table 1). The frequencies of the mutated DegP alleles were 50% and 68.08% in F4 and B2 8 µM populations respectively, and these rose to 84% and 98.61% in the 16 µM populations, respectively (Table 1). The second gene identified in both clones was TGME49_239240, which is annotated in ToxoDB as a serine/threonine protein kinase with similarity to calmodulin-dependent and myosin light chain kinases. Because its function is unknown in *T. gondii*, it is referred to here as artemisinin resistance kinase 1 (Ark1). The mutations in Ark1 occurred at the same site, which resides in the hinge region of the ATP binding pocket, with a different non-conservative substitution in each lineage (Ark1-F4-C274R and Ark1-B2-C274F). Mutations appeared in Ark1 in 70.58% of the reads in the B2 8 µM population but only 4.1% mutant allele reads in the 8 µM population from the F4 line (Table 1). However, in the 16 µM populations, Ark1 mutated alleles reached 90% and 100% abundance in F4 and B2 populations, respectively (Table 1). The frequency of mutations in both genes increased to 100% in the 100µM ART selected populations from both B2 and F4 lines (Table 1).

**Table 1.** Mutations found in candidate genes identified by whole-genome sequencing that were shared in both ART selected lines.

| Line | Gene | Name | Frequency % (ART 8µM) | Frequency % (ART 16µM) | Frequency % (ART 100µM) | Coding region change | Amino acid change |
| --- | --- | --- | --- | --- | --- | --- | --- |
| F4 | TGME49_290840-serine protease | DegP | 50 | 84 | 100 | 2461G>C | Glu821Gln |
| F4 | TGME49_239420-protein kinase | Ark1 | 4.1 | 90 | 100 | 820T>C | Cys274Arg |
| B2 | TGME49_290840-serine protease | DegP | 68.08 | 98.61 | 100 | 2417G>A | Gly806Glu |
| B2 | TGME49_239420-protein kinase | Ark1 | 70.58 | 100 | 100 | 821G>T | Cys274Phe |

### Validation of the DegP and Ark1 mutations in conferring ART tolerance

To establish a functional link between the identified mutations and ART resistance, we used a CRISPR/Cas9 based markerless genome editing strategy to introduce non-synonymous SNPs into the endogenous loci of the wild type parental line (Fig. 2A) (53). Two pairs of mutations were created to simulate mutations identified in the B2 and F4 resistant lines. Each set consisted of clones with single mutants in DegP, single mutants in Ark1, and double mutants of both DegP and Ark1. Initially, we reconstituted the DegP G806E and Ark1 C274F mutations from the F4 line, either alone or together. We then constructed similar mutants based on the DegP E821Q and Ark1 C274R mutations from B2. The point mutations were introduced into a wild type RH Δku80 line expressing firefly luciferase (FLUC), so that we could use a luciferase-based growth assay to monitor growth more precisely. The correct mutations were confirmed by restriction enzyme digestion followed by Sanger sequencing of an amplicon flanking the mutated residue. Surprisingly, introduction of any single- or double-point mutation from either lineage resulted in only a slight increase in EC_50_ ranging from 1.2-1.6-fold (Fig. 2B, 2C)(*SI Appendix*, Table S2) but did not recapitulate their phenotypes shown in Fig. 1C, suggesting that the mutations confer enhanced tolerance to drug. One possible explanation for these results is that additional mutations observed in the original selected populations are required to confer increased ART resistance. However, there were no other mutations found in common between the two lines (*SI Appendix*, Table S1). Additionally, when we analyzed the genomic sequences of B2 and F4 lines grown at 100 µM ART and their respective clones, we also did not detect any additional novel mutations in coding regions that were common to both lineages (*SI Appendix*, Table S1, Datasets S3 and S4).

**Fig 2.**
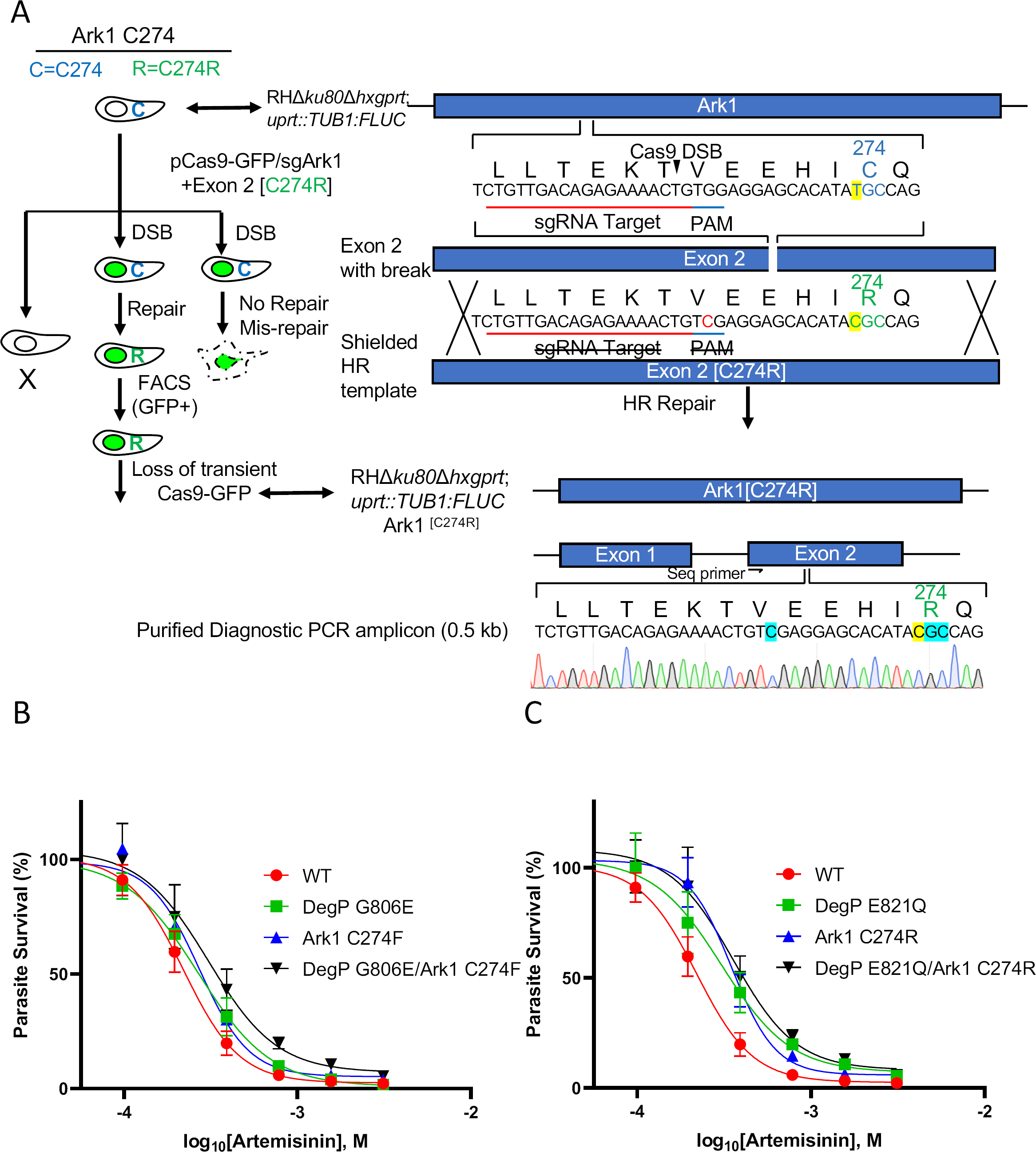
Effect of ART on growth of wild type *T. gondii* and engineered lines containing point mutations in candidate genes. A) Schematic illustration of the markerless CRISPR/Cas9 genome editing strategy used to introduce mutations into the *T. gondii* chromosome for candidate genes. The scheme depicts introduction of the C274R mutation into the Ark1 gene. The wild type allele is C274, while the mutant is C274R. DSB, double-stranded break. HR, homologous recombination. Transformants were isolated by FACS for GFP expression followed by single cell cloning and confirmation of editing by diagnostic PCR and sequencing. B, C) Dose response curves for inhibition of *T. gondii* wild type and point mutant tachyzoites growth in response to increasing concentration of ART. Parasite clones were inoculated in 96-well plates containing HFF cells and growth was monitored by measurement of luciferase activity after 72 hr of incubation with different concentrations ART. Data presented as percent relative light units (% RLU) normalized to growth in naïve (untreated) HFF cells. Shown are three biological replicates each with technical replicates (n=9) ± SE.

To further characterize the role of these mutations in conferring tolerance to ART, we performed a growth competition assay *in vitro* under pressure of ART. In these assays an equal number of wild type and mutant parasites were co-cultured in the presence of 16 µM ART or in vehicle DMSO as a control. When grown in the presence of ART, the mixture of parasites exhibited a slight delay before lysing the monolayer at day 3 vs. the normal day 2 time period. When these parasites were passed onto a fresh HFF monolayer in the presence of ART, they exhibited a growth crash, with variable times of recovery that were dependent on the strains present in culture. Cultures containing DegP mutant parasites were the slowest, with a recovery time of ∼11 days. Cultures containing Ark1 mutants exhibited shorter recovery times ranging between 6 and 8 days. The double mutants in both lineages consistently showed the fastest recovery time ranging between 5 to 6 days.

To analyze the proportion of the wild type vs. mutant strains at the initial inoculation and after the recovery from the ART induced growth crash, genomic DNA was extracted at both time points and the respective loci were PCR amplified and analyzed by Sanger sequencing (Fig. 3A). Remarkably, all single point mutants and their double mutant combinations reproducibly outcompeted the wild type parental strain by the end of the culture period in the presence of ART (Fig. 3B, 3C). Of note, while both DegP point mutants showed a fitness defect when co-cultured with the wild type strain under control conditions, they consistently outcompeted the wild type in the presence of ART (Fig. 3B, 3C). Interestingly, this growth defect was rescued in the double mutant containing both DegP and Ark1 mutations (Fig. 3B, 3C).

**Fig 3.**
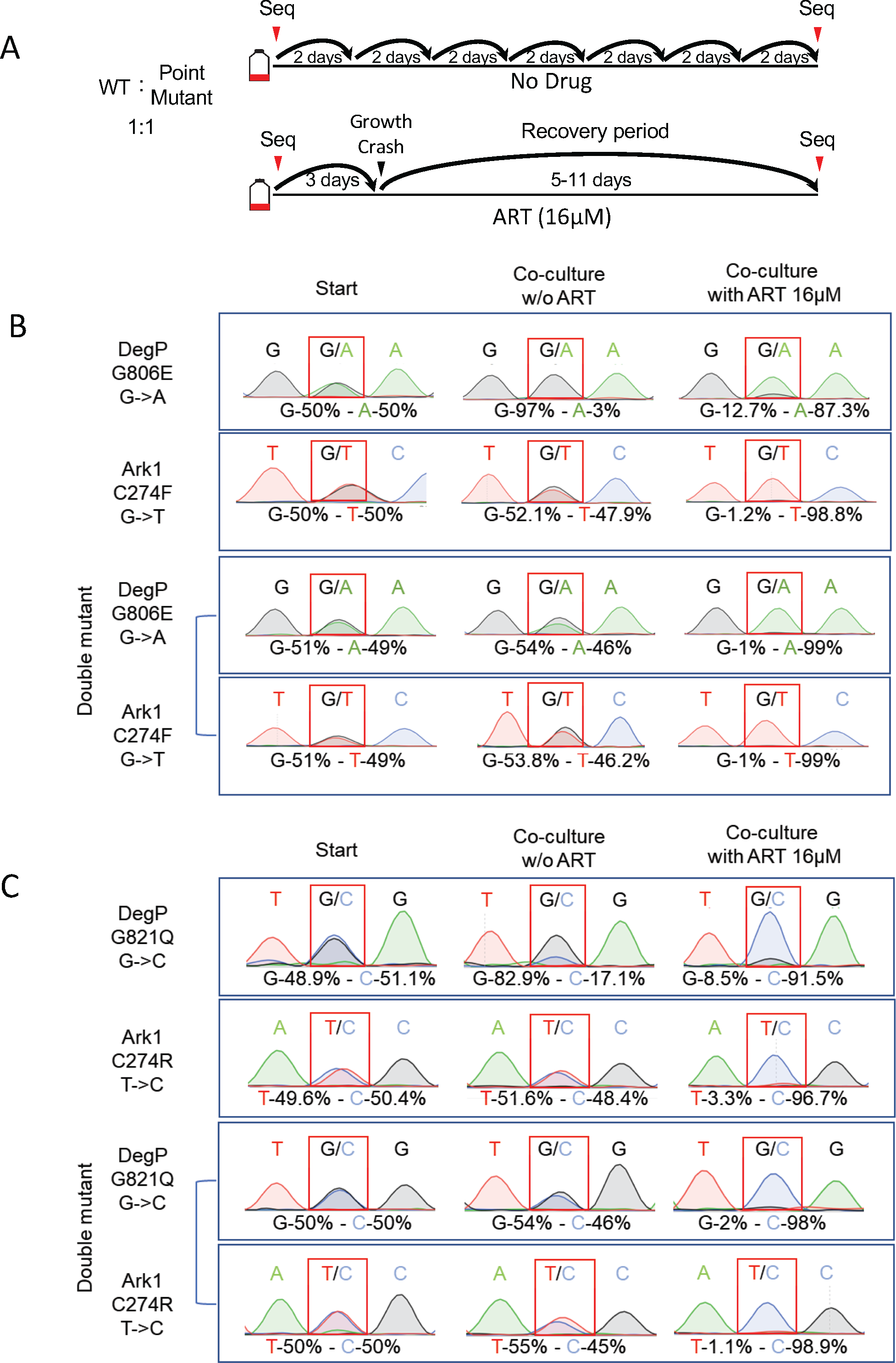
Growth competition assays between wild type and the point mutant strains. **A)** Schematic for growth competition assay. Parental and point-mutant parasites were inoculated at 1:1 ratio (0.5×10^^6^ parasites of each strain) into T25^2^ flasks of confluent HFFs with or without ART 16 μM. After natural egress, parasites were passed by a dilution of 1:10 – 1:20 into a fresh T25^2^ flasks containing confluent HFF cells. Parallel cultures without drug treatment were serially passaged throughout the duration of the experiment. *Toxoplasma* genomic DNA was extracted at the starting time point and after the recovery from the ART induced growth crash. The mutated loci were amplified by PCR and analyzed by Sanger sequencing. DNA sequence traces were used to estimate the relative proportions of sequence variants for wild type and mutant alleles. **B, C**) DNA sequence traces results for analysis of the ratio of wild type and the point mutants. Two different competition assays are shown: (**B**) DegP G806E or Ark1 C274F vs. the combined double mutant and DegP G821Q or (**C**) Ark1 C274R vs. the combined double mutant.

### *T. gondii* ART resistant parasites show amplification of the mitochondrial genome

The resistance to ART in *Plasmodium falciparum* has been previously shown to be associated with increased *pfmdr1* copy-number (54). Hence, we considered the possibility that CNVs within the evolved resistant populations could explain the inability of the single- and double-point mutations to perfectly phenocopy the elevated ART resistance. To examine the genome for CNVs, we mapped the Illumina reads to the whole genome assembly containing the chromosomes of *T. gondii* and examined the read depth as a proxy for copy number. Our initial assessment of CNVs within the resistant populations uncovered four genes (*SI Appendix,* Table S3) with small regions of elevated read depth in the 100µM artemisinin treated clones (*SI Appendix*, Fig. S1). The amplified regions were quite short, consisting of just 200-300 bps, and all four repeats resided in introns of genes with no known association with drug resistance (*SI Appendix,* Table S3). These regions were also elevated in the parental B2 and F4 lines as compared to the rest of the 1X gene, but the read depth increased dramatically in the 100µM artemisinin treated clones (*SI Appendix,* Fig. S1). Interestingly, all four repeats had top BLAST hits annotated as cytochrome b (COB) or cytochrome c oxidase (COX1), genes that are normally encoded in the mitochondria. The current *T. gondii* genome assembly does not include the mitochondrial genome sequence, as its assembly has been complicated by the presence of many short repeats derived from COX1 and COB (55). These genes are normally mitochondrially encoded, but many mtDNA fragments have been amplified and transferred to the nuclear genome of *T. gondii* as scattered imperfect copies that are often flanked by short repeat elements (55). Therefore, we realigned the Illumina reads from the control and the resistant population genomes to the genome assembly, but this time included an independently sequenced portion of the mtDNA spanning COB and COX1 (56) and unique non-assembled supercontigs that likely also contain reads from the mitochondrial genome. This analysis revealed an amplification of contigs that likely constitute the mtDNA, including genes encoding COX1, COX3, and COB (Fig. 4, Dataset S5). The degree of amplification was very dramatic and proportional to the resistance level of the parasites with the ART 100 μM populations and clonal genomes showing the highest level of amplification (Fig. 4, Dataset S5). Analysis of the mitochondrially encoded genes identified only a few SNPs or short indels in a minority of reads (5-10% overall) without any pattern to ART treatment, indicating that the amplifications are not pseudogenes, truncations, or rearrangements, but rather represent increased copies of the intact genome.

**Fig 4.**
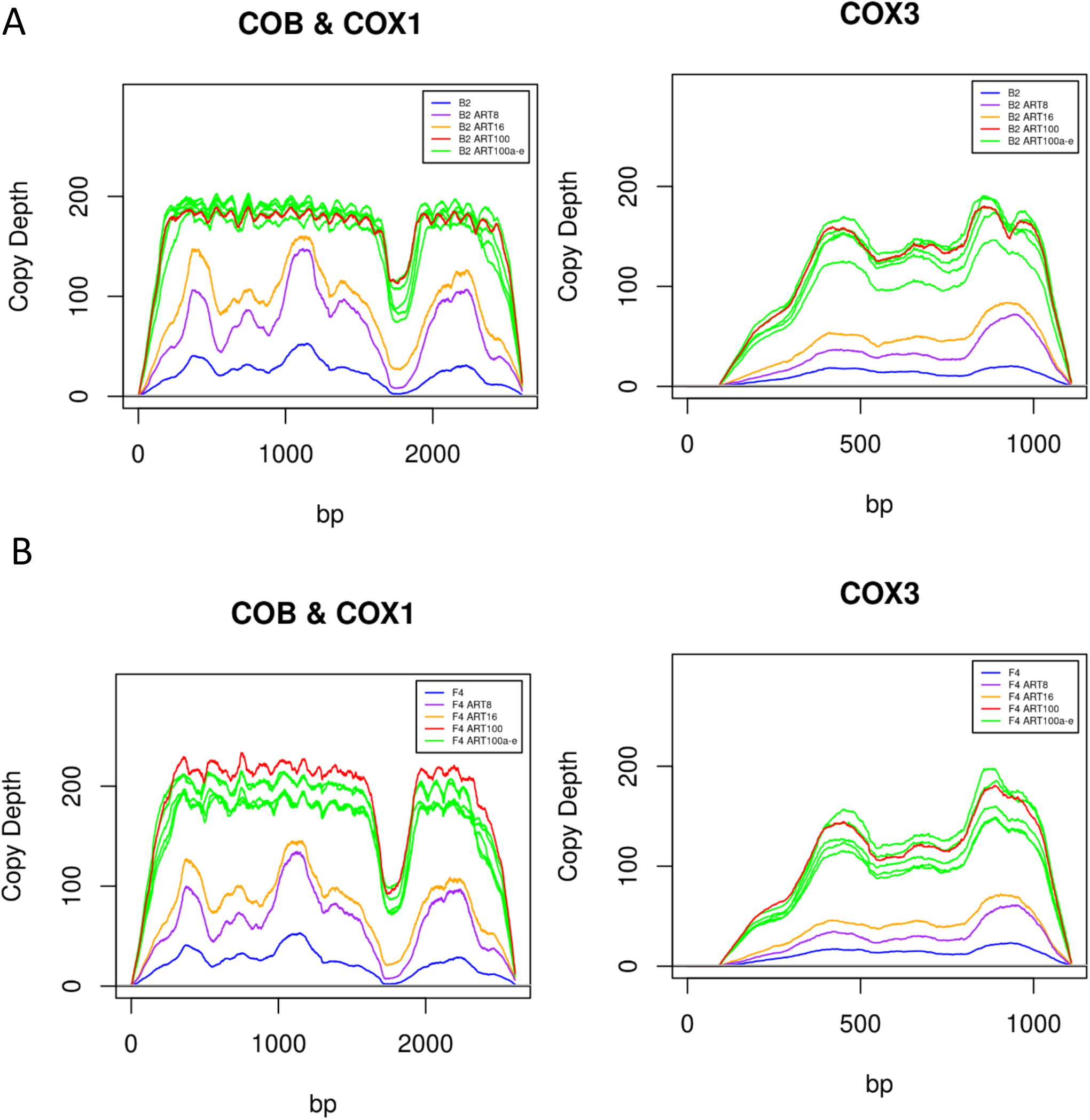
Copy number variation in the mtDNA in ART resistant strains. **A)** Copy number variation for wild type B2 parental and ART-resistant lines derived from this parent. Illumina reads were mapped across mtDNA genes *COB* (GenBank: JX473253.1, 1-1080 bp), *COX1* (GenBank: JX473253.1, 1117-2607 bp), and *COX3* (supercontig KE138885). The B2 lines were either untreated (blue) or treated with 8 µM (purple), 16 µM (orange) or 100 µM (red) artemisinin. The individual 5 clones (a-e) from the 100 µM artemisinin population are shown (green). Copy number estimates are based on the read depth per base pair normalized to 1X across the respective genome. **B)** Copy number variation for wild type F4 parental and ART-resistant lines derived from this parent. Illumina reads were mapped across mtDNA genes *COB* (GenBank: JX473253.1, 1-1080 bp), *COX1* (GenBank: JX473253.1, 1117-2607 bp), and *COX3* (supercontig KE138885). The B2 lines were either untreated (blue) or treated with 8 µM (purple), 16 µM (orange) or 100 µM (red) artemisinin. The individual 5 clones (a-e) from the 100 µM artemisinin population are shown (green). Copy number estimates are based on the read depth per base pair normalized to 1X across the respective genome.

### Artemisinin affects *T. gondii* mitochondrial physiology

Based on the finding that ART-resistant strains showed amplification of the mitochondrial genome, we investigated the effects of ART treatment on *T. gondii* mitochondrial biology. First, we assessed mitochondrial membrane potential (ΔΨm) of the ART-resistant and wild type parental parasites upon treatment with the drug. Intracellular wild type parasites were treated for 24 hr with 8 µM ART or vehicle as a negative control (Fig. 5). Because the ART-resistant parasite populations were kept under constant drug pressure over the course of the evolution experiments, we also examined the effect of ART removal for 24 hr (Fig. 5). Following the various treatments, cultures were stained with MitoTracker, fixed and further stained for a previously characterized mitochondrial protein TgTom40 (57). Intracellular parasites were imaged and examined for changes in mitochondrial localization and membrane potential as indicated by changes of fluorescence intensity and localization of the MitoTracker and TgTom40 signals. The vehicle-treated wild type parasites showed brightly fluorescent mitochondria with a characteristic annular morphology evident in both MitoTracker and TgTom40 staining (Fig. 5A). However, treatment of wild type parasites for 24 hr with 8 µM ART resulted in loss of bright MitoTracker staining, indicative of a perturbed ΔΨm. In contrast, treatment of wild type parasites with ART did not alter the staining pattern or the intensity of TgTom40 (Fig. 5A). We then investigated the ART100 resistant population in the presence of 100 µM ART, conditions under which they grow normally. MitoTracker staining revealed a highly atypical pattern consisting of a few bright foci varying in size with a corresponding reduction in fluorescence intensity in other mitochondrial areas that were still readily stained by TgTom40 (Fig. 5A). Due to the reduced MitoTracker staining in most regions, there was limited overlap with TgTom40 staining (Fig. 5A). Culture of the ART100 population for 24 hr in the absence of drug resulted in an increase in MitoTracker signal throughout the parasite mitochondria with the bright foci remaining intact (Fig. 5A).

**Fig 5.**
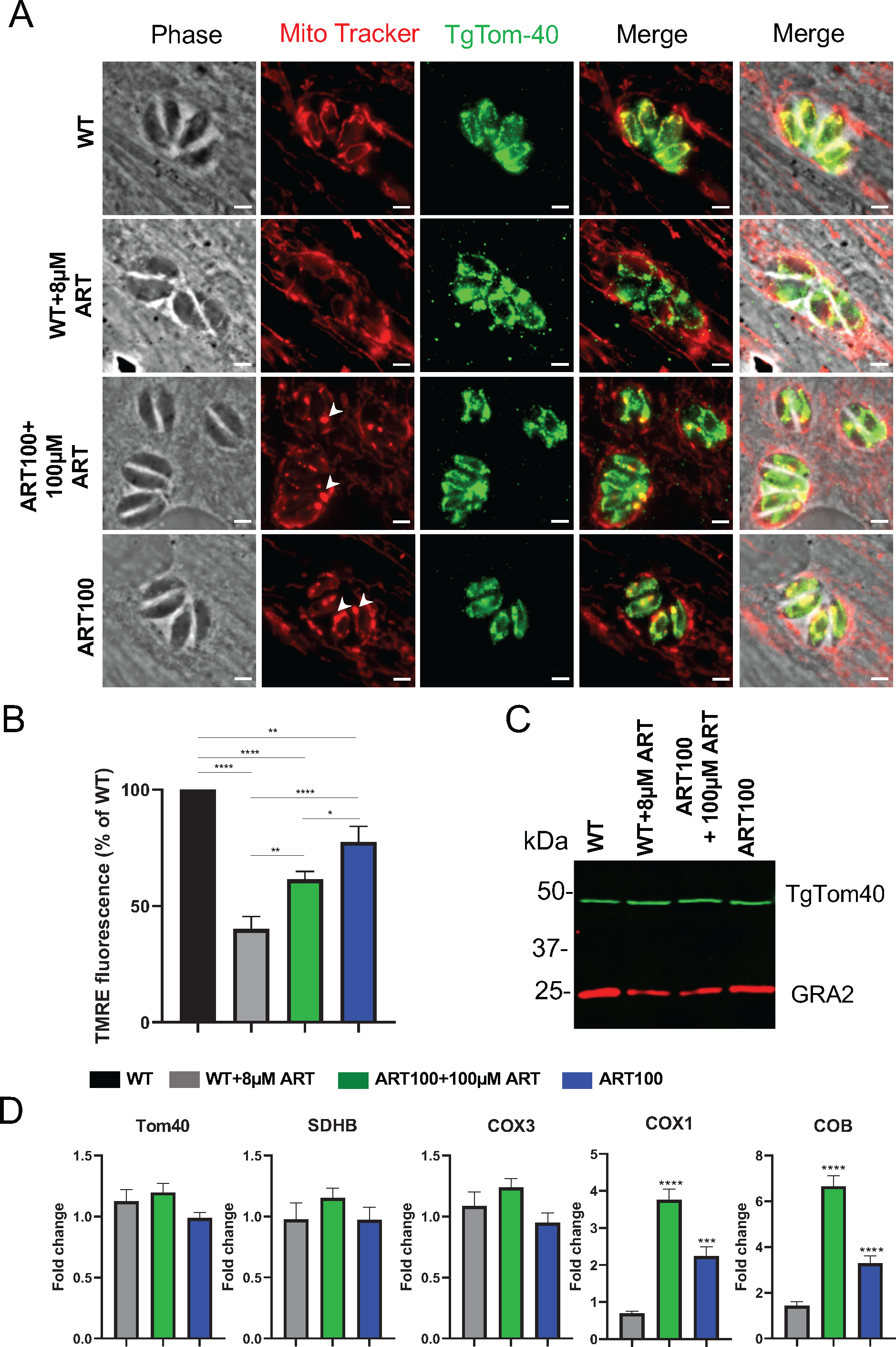
Effects of treatment with ART on *Toxoplasma* mitochondrial morphology and physiology. A) HFF monolayers were infected with wild type or ART100 mutant populations of *T. gondii* and treated with vehicle or ART for 24 hr. Prior to fixation, monolayers were labeled for 30 min at 37°C with 200 nM MitoTracker. Cells were visualized by fluorescence microscopy. Phase, MitoTracker (red), TgTom40 (green), and merged images are shown. Scale bars are 2 µM. B) TMRE assay for mitochondrial membrane potential. HFF monolayers were infected and treated similarly to A). Data are means ± SEM of N = 3 independent biological experiments done with internal duplicates. Statistical analysis was performed using one-way ANOVA with Tukey multiple comparisons test (\*\**P* < 0.001,\*\*\*\**P* < 0.0001) compared to the untreated wild type. C) Immunoblot of lysates from wild type and ART100 parasites treated with ART or vehicle for 24 hr. Blots were probed with rabbit anti-TgTom40 and anti-rabbit IgG IRDye 800RD (green), mouse anti-GRA2 and anti-mouse IgG IRDye 680CW (red). Combined channels of the membrane scan are shown. D) Real-time PCR showing fold induction of mRNA transcripts in wild type or ART100 parasites treated with ART or vehicle for 24 hr. Comparative cycle threshold values were used to evaluate the fold change in transcripts using *ACT1* as an internal transcript control. Data are plotted as fold change ± SEM compared with wild type vehicle treated cells from at least 3 independent experiments per gene. Statistical analysis was performed using one-way ANOVA with Dunnett multiple comparisons test (\*\*\*\**P* < 0.0001) compared to the untreated wild type.

To quantify the effects of ART treatment on ΔΨm, we utilized TMRE (tetramethylrhodamine, ethyl ester) to label active mitochondria. TMRE is a cell permeant, positively charged, red-orange dye that readily accumulates in active mitochondria due to their relative negative charge. Following infection of HFF cells, the monolayers were labeled with TMRE, the parasites were mechanically lysed, and fluorescence levels were read spectrophotometrically in a microtiter plate. Consistent with the observations made with MitoTracker, ART treatment of wild type parasites reduced parasite ΔΨm by 60% (Fig. 5B). Furthermore, the ART100 population grown in the presence of 100uM ART showed significantly higher ΔΨm values compared to those of ART-treated wild type parasites (Fig. 5B). Removal of the drug from the ART100 population exposed to ART for 24 hr lead to a significant increase in ΔΨm, although it did not fully return to wild type levels (Fig 5B).

We next asked whether the amplification of mitochondrial genome leads to a concomitant increase in nuclear encoded mitochondrial components at both the protein and mRNA levels. To answer this question, protein levels of TgTom40 in wild type and mutant strains were examined by Western blot. Neither the addition of ART to wild type parasites, nor removal of the drug from the ART100-resistant population led to a change in TgTom40 levels when compared to that of the untreated wild type cells (Fig. 5C). Nuclear genes that encode proteins imported into the mitochondrion (i.e. *TOM40* and Succinate dehydrogenase [ubiquinone] iron-sulfur subunit succinate dehydrogenase *SDHB*) showed similar transcript levels in wild type parasites upon addition of ART, and in ART100 populations in the presence or absence of drug (Fig. 5D). In contrast, the ART100 population showed significantly higher transcript levels for the mitochondrially encoded *COX1* and *COB* genes compared to wild type parasites (Fig. 5D). Transcript levels for these genes were also responsive to ART removal, resulting in a significant drop (Fig. 5D) which was consistent with the change in ΔΨm in ART100 population grown without the drug Fig. 5B).

Finally, we characterized ART effects on the ultrastructure of mitochondria by transmission electron microscopy (TEM). Mitochondria of wild type parasites treated with vehicle showed a typical dense matrix with multiple clearly defined cristae (Fig. 6A). Treatment of wild type parasites with 8 µM ART for 24 hr resulted in expansion of the matrix, loss of density, and a reduced number of cristae (Fig. 6B). The mitochondrial ultrastructure of the ART100 resistant population grown in presence of 100µM ART presented further reduction in matrix density with only few visible cristae (Fig. 6C). Removal of the drug from the ART100 population for 24 hr led to partial increase in matrix density and restoration in cristae numbers (Fig. 6D).

**Fig 6.**
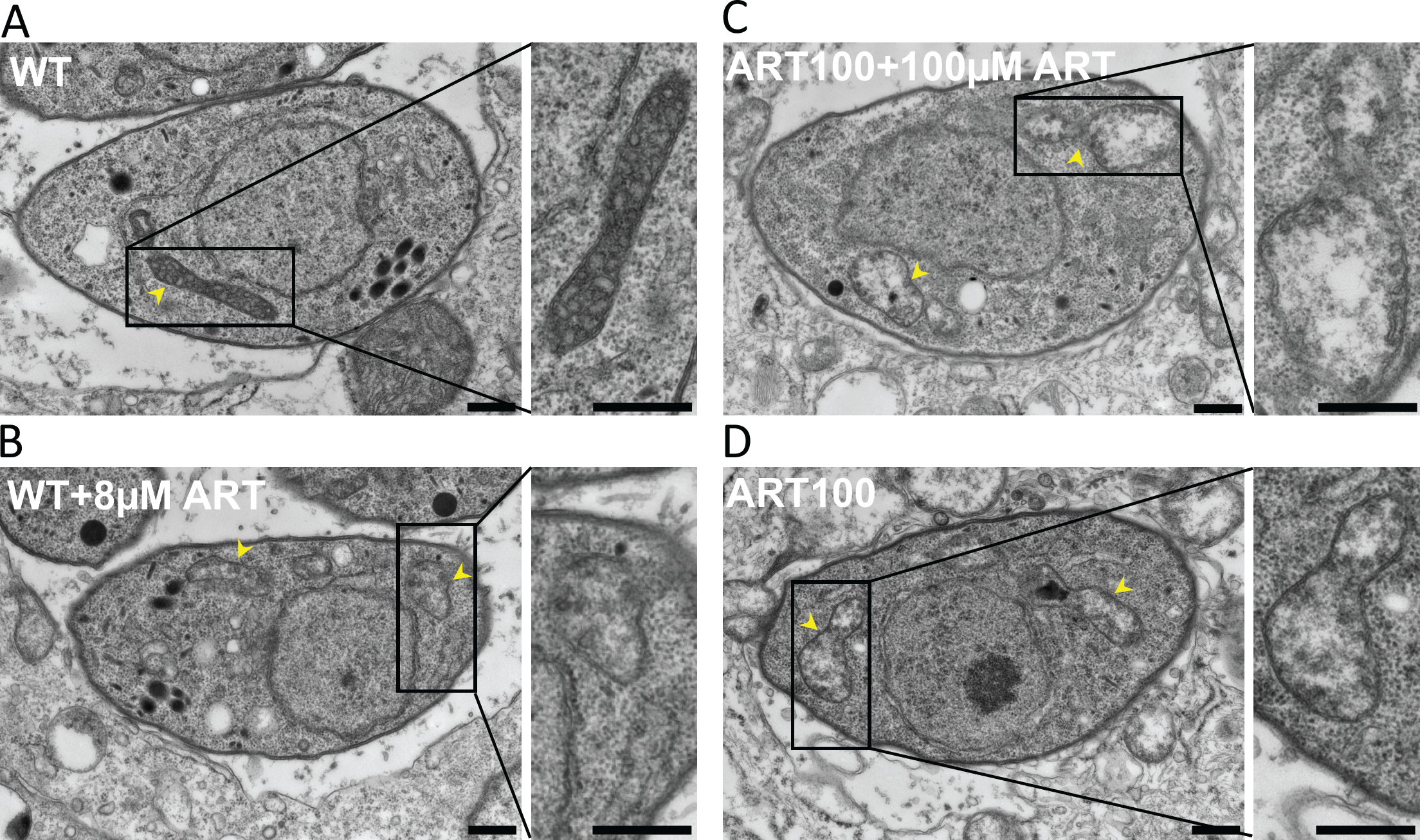
ART effects on *T. gondii* mitochondria ultrastructure. A) Electron micrographs of wild type parasites treated with vehicle (24 hr) B) Electron micrographs of wild type parasites treated with 8 µM ART for 24 hr C) Electron micrographs of ART100 resistant population treated with 100 µM ART D) Electron micrographs of ART100 resistant population grown without ART for 24hr. Arrowheads indicate parasite mitochondria. Scale bars are 500 nm.

## Discussion

We took advantage of flexible *in vitro* culture systems and genetic tools available in *T. gondii* to evolve lines that were resistant to growth-inhibiting effects of ART in order to define potential mechanisms of action. In parallel, we determined the background mutation rate of clones generated after long-term passage in the absence of ART selection. Passage in ART led to enhanced resistance at the population level and whole genome sequencing in the ART-resistant lines identified several non-synonymous SNPs in the coding sequences of a serine protease homologous to DegP and a serine/threonine kinase named Ark1. We engineered these point mutations into a wild type background using CRISPR/Cas9 and confirmed that they confer a competitive advantage in the presence of ART, although they did not alter the EC_50_ value for growth inhibition. Among the ART-resistant lines, we also observed upregulation of heme-containing cytochromes encoded in the mitochondrial genome, consistent with previous studies suggesting a mitochondrial target for ART. Collectively our studies define new mechanisms of mutation that result in ART resistance, including point mutations in specific genes that have not been seen previously, and alterations in mitochondrial function.

Analysis of the frequency of mutations in clones of *T. gondii* that were grown under standard laboratory conditions led to several surprising findings. First, the rate of apparent mutation in *T. gondii* was relatively low at ∼ 5.8 ×10^-11^ / mutations / bp / generation. This rate is ∼ 30 fold lower than that estimated for *P. falciparum* (1.7 ×10^-9^) (58), but somewhat closer to rates described in Baker’s yeast (3.3 ×10^-10^) (59), or *S. pombe* (2.0 ×10^-10^) (60). Second, we observed a high level of nonsynonymous (dN) relative to synonymous (dS) mutations in coding regions. In the absence of selection, this ratio is expected to be ∼ 1, indicating that strong selection was operating under standard growth conditions. As many nonsynonymous mutations have seemingly been lost from the clones, the true mutation rate is likely to be higher. Previous studies examining SNPs between different lineages of the RH strain that were passaged for an unknown number of doublings in different laboratories also reported elevated dN/dS ratios that ranged from 2 - 10 (61). Studies conducted on the model organism *E. coli* have shown that strong selective pressures exist even under homogeneous conditions and that mutations fluctuate widely over time in frequency but often reach fixation (62). Such mutations also show strong dN/dS skewing and often contribute to fitness advantages (62). In the present study, we did not evaluate the fitness of the mutations that reached fixation in non-selected lines, although the high dN/dS ratio is more consistent with improved growth characteristics rather than potential bottlenecks during passage. Our passaging procedures are highly similar to what most laboratories use for maintenance of *T. gondii* cultures *in vitro*, suggesting this could be a widespread phenomenon.

Following culturing at elevated levels of ART, we obtained two parallel populations with increased resistance to ART as evidenced by shifted EC_50_ values in an *in vitro* growth assay. Whole genome sequencing of these lines revealed a number of mutations in each lineage, but two genes stood out as having a high frequency of SNPs in the population and clones from both lines. Since changes in these two genes were most clearly associated with increased resistance, we chose to examine them further. We reintroduced the point mutations seen in the resistant lines, either alone or in combination, into wild type background using CRISPR/Cas9 to edit the genome. Somewhat surprisingly, introduction of these point mutations did not shift the EC_50_ to a similar extent as seen in the original uncloned populations. Hence, the presence of these mutations alone appears to confer tolerance, but not outright resistance to ART. Although the observed mutations do not correspond to those previously identified in *P. falciparum*, the phenotype is reminiscent of the phenotype seen in the RSA where “resistant” lines show improved survival but do not exhibit a shift in the EC_50_ (29). The reasons for the difference in EC_50_ shift between the engineered clones and the original lines are unclear but might relate to additional background mutations in each of the evolved lineages, or due to alterations in the mitochondrial genome, as discussed below. Regardless, the point mutations did enhance fitness in a competition assay with wild type parasites grown in the presence of ART. Each of the single and double mutants outgrew the wild type cells within the first several passages, despite a lag in the initial expansion. This enhanced fitness was not due to improved growth overall as the DegP mutants showed a growth disadvantage, while the Ark1 mutants were equal to wild type in the absence of ART. Interestingly, the Ark1 mutants rescued the partial growth defect of the DegP mutants in the absence of ART. However, this was not the sole reason for expansion of this allele as the single Ark1 mutations also conferred a competitive advantage vs. wildtype parasites in the presence of ART.

The exact mechanisms by which point mutations in these genes confer enhanced tolerance to ART in *T. gondii* are unclear but neither of these two genes have been implicated in previous studies on ART resistance in other systems. The mutations seen in *T. gondii* Ark1 occur in the hinge region of the kinase, a critical spot for ATP binding, suggesting they might affect catalytic activity. The substrates of Ark1 are presently unknown and its role in the biology of the parasite has not previously been explored, although it has a fitness score in a genome-wide CRISPR screen that would suggest it is essential (−3.77) (63). Similarly, the functional consequences of the DegP mutations in conferring tolerance to ART in *T. gondii* are unclear from the present studies. DegP is a member of the High Temperature Requirement A family (HtrA) and is a periplasmic protease in *E. coli* (64). In bacteria, DegP is involved in protective stress responses to aid survival at high temperatures (64), a phenotype also reported in yeast (65). Htr orthologs in eukaryotes are typically found in the mitochondria where they function in quality control (66). Given the role of DegP in stress responses, it may play a similar role to K13 in ART resistance in malaria, where mutations in K13 propeller domains have been linked to altered stress responses (32). By analogy, the mutations in the PDZ domains seen in ART resistant *T. gondii* may affect chaperone activity and/or interaction with its substrates and in the process relieve stress induced by treatment with ART.

Consistent with previous studies in yeast (43) and tumor cells (45, 46), our findings support the mitochondrion as a target of artemisinin. We observed both a decrease in membrane potential and altered morphology in wild type parasites treated with ART and a stable decrease in membrane potential and altered morphology in resistant populations, suggesting these changes are a long-term adaptation to the presence of drug. Initial studies in yeast suggested that the electron transport chain (ETC) was responsible for activation of the endoperoxide moiety in artemisinins (27, 43), a functional group that is important for its toxic activity. However, mammalian cell petite mutants, which lack a functional ETC, show no deficiency in formation of cytotoxic radicals and instead chemical modulators support a role for heme in activating the endoperoxide group (46). Nonetheless, such ETC-deficient mammalian cells are more resistant to artemisinins, supporting a role for this pathway through generation of reactive oxygen species and induction of apoptosis (46). The ART100 population of *T. gondii* mutants studied here demonstrated reduced ETC activity in both the presence and absence of drug, a feature that would be expected to contribute to enhanced resistance. This suggests that the amplification of the mitochondrial genome and concomitant enhanced transcription of several subunits encoded there creates an imbalance in the components of the ETC, thus impairing its function. Future studies are required to resolve the roles of heme vs. the ETC in activating ART in *T. gondii*, as well as identifying the targets and pathways that result in cytotoxicity.

Our findings provide support for several previously suggested mechanisms of ART resistance and also make several new predictions about the frequency and consequences of mutation in *T. gondii*. First, we identify new point mutants that can confer enhanced survival in the presence of ART without shifting the EC_50_ in a classical resistance phenotype. Although the target genes differ, this finding mirrors what is seen in *P. falciparum* where K13 mutations are associated with enhanced survival in the presence of drug. Second, similar to *P. falciparum*, the mechanisms of resistance to ART in *T. gondii* appear to be multigenic and may have a link to altered stress responses. Third, ART may inhibit mitochondrial function in *T. gondii* and alterations in targets there may thus lead to resistance, as previously suggested by studies done in yeast and tumor cells. Fourth, although the low intrinsic mutation rate in *T. gondii* may limit the potential for drug resistance to arise in the clinic, our finding that mutational drift occurs readily in the absence of purposefully applied selection predicts that mutations that may alter phenotypes are likely to accumulate during normal laboratory passage. Collectively, these processes may also drive evolution and affect the occurrence of drug resistance and other biological traits *in vivo*.

## Materials and Methods

### Reagents and antibodies

All chemicals were obtained from Sigma (St. Louis, MO) or Thermo Fisher Scientific (Waltham, MA) unless otherwise stated. MitoTracker™ Red CMXRos was purchased from Life Technologies (Carlsbad, CA). Secondary antibodies used for immunofluorescence were conjugated to Alexa Fluor 488, Alexa Fluor 568, or Alexa Fluor 350 and were purchased from Invitrogen (Carlsbad, CA). The following primary antibodies were used: rabbit polyclonal anti-TgTom40 antibody (a generous gift of Giel Van Dooren, Australian National University), (57)), and mouse monoclonal GRA2 (67). For ART preparations, Artemisinin (Sigma) was prepared as 250 mM stock in 100% DMSO and stored at −80 °C until use. ART was further diluted in DMEM medium to achieve the required final concentration with DMSO final concentration of 0.1%.

### Parasite and Cell Culture

The type I *T. gondii* strain RH-88, transgenic lines reported previously RHΔ*hxgprt*Δ*ku80* (68), or developed here are listed in (*SI Appendix*, Table S4). *T. gondii* tachyzoites were maintained in human foreskin fibroblast (HFF) monolayers cultured at 37°C, 5% CO_2_ in D10 medium [Dulbecco’s modified Eagle’s medium (Invitrogen)] supplemented with 10% HyClone fetal bovine serum (GE Healthcare Life Sciences), 10 μg/mL gentamicin (Thermo Fisher Scientific), 10 mM glutamine (Thermo Fisher Scientific)]. HFF cells were grown as confluent monolayers in T25^2^ flasks and inoculated with ∼2×10^6^ parasites that expanded by ∼ 10 fold over 2-3 days of culture. Following natural egress, the cultures were split by 1:10 or 1:20 dilution of the growing culture to a fresh uninfected HFF monolayer growing in a T25^2^ flask. In order to harvest parasites for the assays described below, parasites were allowed to egress naturally and tachyzoites were purified by filtration through 3 micron polycarbonate filters (Whatman) in Hank’s balanced salt solution supplemented with 0.1 mM EGTA and 10 mm HEPES, pH 7.4, followed by centrifugation at 400 *g*. Clones were isolated by limited dilution and outgrowth on monolayers of HFF cells grown in 96 well plates. All strains and host cell lines were determined to be mycoplasma negative using the e-Myco plus kit (Intron Biotechnology).

### Parasite Transfection

Following natural egress, freshly harvested parasites were transfected with plasmids, using protocols previously described (69). In brief, ∼2 x 10^7^ extracellular parasites were resuspended in 370 μl cytomix buffer were mixed with ∼ 30 μl purified plasmid and/or amplicon DNA in a 4 mm gap BTX cuvette and electroporated using a BTX ECM 830 electroporator (Harvard Apparatus) using the following parameters: 1,700 V, 176 μs pulse length, 2 pulses, 100 msec interval between pulses. Transgenic parasites were isolated using a FACSAria II (BD Biosciences) fluorescence-activated cell sorter (FACS) on the basis of Cas9-GFP fluorescence or by outgrowth under selection with pyrimethamine (3 µM) or 5-fluorodeoxyuracil (10 µM), as needed. Stable clones were isolated by limiting dilution on HFF monolayers grown in 96-well plates.

### Plasmid Construction and Genome Editing

All CRISPR/Cas9 plasmids used in this study were derived from the single guide RNA (sgRNA) plasmid pSAG1:CAS9-GFP, U6:sgUPRT (70) by Q5 site-directed mutagenesis (New England Biolabs) to alter the 20 nt single guide RNA (sgRNA) sequence, as described previously (71). Primers for plasmids are listed in *SI Appendix* Table S5. Separate sgRNA plasmids were made for editing mutations in DegP (i.e. pSAG1:CAS9-GFP, U6:sgDegP ^G806E^; pSAG1:CAS9-GFP, U6:sgDegP ^[G821Q]^;) and Ark1 (i.e. pSAG1:CAS9-GFP, U6:sgArk1).

### Generation of Firefly Luciferase (FLUC) Tagged Strain – Rh*luc*

To tag RH Δ*KU80* strain with FLUC, the reporter plasmid pUPRT::Floxed DHFR-TS*,TUB1:Firefly Luciferase plasmid (72) was targeted to the *UPRT* gene in *T. gondii* by co-transfection with the CRISPR plasmid pSAG1:CAS9, U6:sgUPRT. Parasites were sequentially selected in pyrimethamine (1.0 μM) followed by 5-fluorodeoxyracil (FUDR, 10 μM) and independent clones isolated by limiting dilution.

### Generation of point mutant parasites in endogenous genes

To introduce point mutations into endogenous genes, we used a markerless genome editing strategy. RH*luc* parasites were co-transfected with a dual-functional Cas9-green fluorescent protein (GFP) / sgRNA plasmid designed to cut within the vicinity of the codon of interest and a Cas9-shielded homology donor amplicon. This amplicon provides a repair template (∼500 bp) containing the point mutation of interest along with a silent mutation that would either introduce a novel restriction site or eliminate an existing one (*SI Appendix* Table S6). Following transfection, parasites transiently expressing Cas9-GFP were isolated by FACS, as described above. Single clones were isolated from the FACS-sorted population by limiting dilution on HFF monolayers in 96 well plates. Following isolation of clones, a 500 bp fragment surrounding the mutant allele was amplified by PCR using primers described in *SI Appendix* Table S4 followed by diagnostic restriction enzyme digestion that would distinguish the wild type and edited alleles. Clones that were positive by restriction enzyme digestion were expanded and further confirmed sequenced by the Sanger.

### Immunofluorescence microscopy

For IF microscopy of intracellular *T. gondii*, HFF monolayers grown on glass coverslips were fixed in 4% formaldehyde for 15 min at room temperature, blocked using 10% normal goat serum (NGS) with 0.1% Triton X-100 in PBS for 15 min, and incubated with primary antibodies in 10% NGS in PBS for 90 min. Cells were washed three times with PBS and incubated with Alexa-conjugated secondary antibodies and Hoechst stain (100 ng/ml) to stain nuclei (Life Technologies) for 30 min. Cells were then washed three times with PBS followed by mounting using ProLong Diamond antifade reagent (Life Technologies) to minimize bleaching during microscopy image acquisition. Standard wide-field images were captured and analyzed with a 63 × or 100 × oil objective on an Axioskop 2 MOT Plus wide-field fluorescence microscope (Carl Zeiss, Inc.) running AxioVision LE64 software (Carl Zeiss, Inc.).For staining with MitoTracker™ Red CMXRos (Thermo), medium was replaced with prewarmed DMEM containing MitoTracker at a concentration of 200 nM. After 30 min of incubation at 37°C, cells were washed and then fixed. Parasites were labeled with rabbit 1:2,000 anti-TgTom-40 as primary antibody followed by 1:2,000 anti-rabbit IgG-Alexa 488 as secondary antibody.

### TMRE Mitochondrial Membrane Potential Assay

To quantify the *Toxoplasma* mitochondrial membrane potential HFFs were infected at MOI 5:1, 24hr post infection the media was changed for DMEM media containing TMRE at a concentration of 500 nM. After 30 min of incubation at 37°C, cells were washed 3 times with IC buffer (142 mM KCl, 5 mM NaCl, 1 mM MgCl2, 5.6 mM D-glucose, 2 mM EGTA, 25 mM HEPES, pH 7.4) and the parasites were mechanically released by scraping and syringe passage. The parasites were centrifuged followed by resuspension in IC buffer to a final concentration of 15×10^6^ / ml. Parasites were transferred in triplicates into to 96-well µCLEAR black plates (Greiner Bio International) at 100µl per well. The wells were analyzed at Ex/Em = 549/575 nm with Cytation3 (Biotek). Mechanically released wild type parasites without TMRE treatment were used for the estimation of the background signal.

### Transmission electron microscopy

For ultrastructural analyses, samples were fixed in 2% paraformaldehyde-2.5% glutaraldehyde (Polysciences Inc., Warrington, PA) in 100 mM sodium cacodylate buffer, pH 7.2, for 2 hr at room temperature and then overnight at 4°C. Samples were washed in sodium cacodylate buffer at room temperature and postinfection fixed in 1% osmium tetroxide (Polysciences Inc.) for 1 hr. Samples were then rinsed extensively in distilled water (dH_2_O) prior to en bloc staining with 1% aqueous uranyl acetate (Ted Pella, Redding, CA) for 1 hr. Following several rinses in dH_2_O, samples were dehydrated in a graded series of ethanol and embedded in Eponate 12 resin (Ted Pella Inc.). Sections of 95 nm were cut with a Leica Ultracut UCT ultramicrotome (Leica Microsystems Inc., Bannockburn, IL), stained with uranyl acetate and lead citrate, and viewed on a JEOL 1200 EX transmission electron microscope (JEOL USA Inc., Peabody, MA) equipped with an AMT 8-megapixel digital camera and AMT Image Capture Engine V602 software (Advanced Microscopy Techniques, Woburn, MA) as part of the Microbiology Imaging Facility, Washington University in St. Louis.

### Western blotting

Cell lysates of HFFs were prepared using CellLytic M (Sigma) mixed with Complete Mini protease inhibitor cocktail (Roche). Cell supernatants were collected after centrifugation at 6,000 × g for 10 min at room temperature (RT) to avoid cell debris. Total protein was measured in each sample using the bicinchoninic acid (BCA) protein assay kit (Pierce, Thermo Fisher Scientific). Samples were boiled at 95°C for 15 min in Laemmli buffer containing 100 mM dithiothreitol (DTT). Samples were separated using SDS-PAGE and transferred onto a nitrocellulose membrane. The membrane was blocked in a 1:1 mixture of Odyssey blocking buffer (OBB; Li-Cor Biosciences) and PBS overnight at 4°C. The membrane was incubated with rabbit polyclonal anti-TgTom40 and mouse anti-GRA2 at 1:2,000 and 1:1,000 respectively for 2 hr at RT in a 1:1 mixture of Odyssey blocking buffer and PBS with 0.1% Tween 20 (PBS-Tween). The blot was washed three times for 5 min each with PBS-Tween and incubated with anti-rabbit IgG IR800 and anti-mouse IgG IR680 at 1:15,000 for 2 hr at RT in a 1:1 mixture of Odyssey blocking buffer and PBS-Tween. The blot was washed three times for 5 min each with PBS-Tween followed by infrared imaging on a Li-Cor Odyssey imaging system.

### Real-time PCR

Samples were lysed, and RNA was extracted using the Qiagen RNeasy minikit per the manufacturer’s instructions. cDNA was prepared using the Bio-Rad iScript cDNA synthesis kit per the manufacturer’s instructions. Real-time PCR of all the genes was performed using Clontech SYBR Advantage qPCR premix per the manufacturer’s instructions. Data acquisition was done in QuantStudio3 (Applied Biosystems) and analyzed in QuantStudio design and analysis software (Applied Biosystems). Primers are listed in Table S1. Comparative threshold cycle (CT) values were used to evaluate fold change in transcripts using actin as an internal transcript control.

### *In Vitro* Growth Assays

#### Lactate dehydrogenase (LDH) release

To assess ART sensitivity of the original resistant mutant clones, growth was monitored over a 10-point dose–response curve based on LDH release from the HFF monolayer. Briefly, 5 × 10^5^ parasites (100 μL/well) were added to HFF cells grown in a 96-well plate that contained 100 μL of 2x ART concentration (to achieve 1x final ART concentration in 200 μL total well volume containing 0.1% DMSO) and allowed to replicate for 48 hr. Parasite growth was quantified by measuring lactate dehydrogenase (LDH) release from host cells as a consequence of rupture, as described previously (73). Supernatants were collected after 48 hr and LDH measured using the CytoTox 96 assay (Promega) according to the manufacturer’s instructions. Values were normalized to total lysis (100%) and uninfected (0%).

#### Luciferase assay

To assess the ART sensitivity of the engineered point mutant strains, growth was monitored over a 10-point dose-response curve using luciferase activity. Briefly, 2.5 x 10^3^ luciferase expressing parasites (100 μL/well) were added to HFF cells grown in a 96-well plate that contained 100 μL of 2x ART concentration (to achieve 1x final ART concentration in 200 μL total well volume containing 0.1% DMSO) and allowed to replicate for 72 hr prior to measuring luciferase activity. Briefly, culture medium was aspirated and replaced with 40 μL of 1x Passive Lysis buffer (1x PLB, Promega, E1531) and incubated for 5 min at room temperature (RT). Luciferase activity was measured on a Cytation 3 (BioTek) multimode plate imager using the following protocol: inject 100 μL of Luciferase Assay Reagent (LAR), shake 1 s, and read 10 s post injection.

Data were analyzed using Prism (GraphPad) to determine EC_50_ values by plotting normalized, log-transformed data (x axis), using non-linear regression analysis as a sigmoidal dose–response curve with variable slope. The EC_50_ data are presented as the average of three biological replicates (i.e., separate EC_50_ titrations) each with three technical replicates (i.e., separate wells).

### Growth Competition Assays

Parental and point-mutant parasites were inoculated at 1:1 ratio (0.5×10^6^ parasites of each line) into T25^2^ flasks of confluent HFFs with or without ART (16 μM). Following growth for several days and natural egress, cultures were passed into a fresh T25^2^ flask at a dilution of 1:10 or 1:20. Parasite cultures growing in the presence of ART exhibited growth crash upon second passage. Parallel cultures of the mixture of wild type and mutant lines were serially passaged with no drug treatment. Parasite genomic DNA was extracted at the starting time point and after the recovery from the ART induced growth crash. The target loci were amplified by PCR and analyzed by Sanger sequencing. QSVanalyser (74) was used for the analysis of DNA sequence traces for estimation of the relative proportions of the wild type and mutant variants.

### Whole-genome sequencing

Parasite DNA was extracted from *in vitro* cultures using QIAamp DNA blood kits (Qiagen). Integrity of genomic DNA was determined on an Agilent Tapestation. An aliquot of 1 μg of gDNA as determined by Qubit assay was sonicated to an average size of 175 bp. Fragments were blunt ended using T4 DNA Polymerase, Klenow DNA Polymerase, and T4 polynucleotide kinase. Then, the fragments were modified to contain an “A” base to 3’ end with Klenow (3′→ 5′ exo-), and ligated to Illumina’s sequencing adapters with T4 DNA ligase. Half of the ligated fragments underwent amplification for 8 cycles incorporating a unique indexing sequence tag with VeraSeq DNA Polymerase. All enzymes were purchased from QIAGEN. The resulting libraries were normalized and sequenced using the Illumina HiSeq-3000 as paired end reads extending 150 bases from both ends of the fragments. Illumina’s bcl2fastq utility was used to demultiplex the samples.

### SNP analysis

Reads were aligned to the *T. gondii* ME49 reference genome (ToxoDB release 41) using BWA-mem (75) and further processed using Picard Tools (http://broadinstitute.github.io/picard/). SNVs and INDELs were called using GATK HaplotypeCaller, then filtered based on GATK recommendations (76). Briefly, SNV calls were retained if they met the following criteria: ReadPosRankSum >8.0 or <-8.0, QUAL<500, Quality by Depth (QD) <2.0, Mapping Quality Rank Sum <-12.5, and filtered depth (DP) <7. Indels were retained if they met the following criteria: ReadPosRankSum <-20, QUAL<500, QD<2, and DP<7. Mutations were annotated with a custom SnpEff (77) database built from a GFF file. Variants were further filtered by removing mutations that were present in both the parent strain and evolved strains, such that mutations were only retained if they arose during the course of long-term culturing for neutral mutation rate experiments or the drug selection process for artemisinin-pressured lines. Manual inspection of the resulting SNPs was used to remove polymorphisms where the read depth for a majority of clones was below the cutoffs stated above, since correct assignment of genotypes was ambiguous in many of these cases.

### Statistical analysis of mutation rate by whole genome sequencing

The mutation rate was calculated based on the number of SNPs identified in each of the subclones from the unselected 5D and 12B lines sampled at 182 or 365 days. For these calculations, we assumed that the growth rate and mutation rate are constant across the experiment. We also assumed that the observed mutations (SNPs) are an accurate measure of the mutation rate across the experiment. The equation used for calculating the mutation rate μ was:

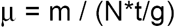

where m = the number of SNPs, N = the genome size of 65,464,221 bp (47), t = time of growth in days and g = the number of doubling per day (assumed to be 4 here (50)). Identical mutation that were observed in more than one clone were only counted once (as designated in *Datasets* S1, S2). We calculated the average mutation rate for the 365 day clones from the individual clones for the two separate lines.

We also estimated the mutation rate by treating the whole genome sequencing results for all unselected clones from the 56D and 12B series as a fluctuation analysis, scoring the number of SNPs (mutations) for each of the clones over 728 doublings equal to 182 days of culture (for clones sampled at 365 days the number of SNPs was halved to allow direct comparison), under the assumption that the mutants have equal fitness to the wild type. Mutation rates were calculated using bz-rates (http://www.lcqb.upmc.fr/bzrates) (78) based on equations that have been described previously (79).

### Copy number variation (CNV) analysis

Estimation of CNV was calculated as previously described (47). Briefly, for each sample, FASTQ files were aligned to a curated *Toxoplasma gondii* ME49 reference genome (http::/ToxoDB.org, v.42), see below, using Bowtie2 version 2.3.4.3 with the –end-to-end option (80) and SAM files were converted to BAM format using samtools version 1.9 (81). Average read depth and standard deviation for each gene in the *T. gondii* genome was obtained using samtools mpileup option across the genomic start:stop positions for each annotated CDS. The genomic 1X mean read depth for each sample was calculated as the average read depth across all base pairs within genomic start:stop positions for CDSs with average read depth between the first and third quartiles off all genes, as determined using the quartile function in R. Copy depth is either the gene average or bp level read depth normalized to the 1X genomic mean for that sample. For both analyses described below, CNV for control genes TGME49_308090 (ROP5 – 6-7 copies), TGME49_212740 (AAH2 – 2 copies), and TGME49_271580 (TgFAMD – 2 copies) match previous estimates of CNVs in the type I RH strain (47)(highlighted yellow in Dataset S5).

Initially, the CNV analysis was performed by aligning to a curated *T. gondii* ME49 genome that included the 14 chromosomes and the apicoplast genome (KE138841) without the unassigned KE supercontigs, named Tg14+api. Most unassigned KE supercontigs are short (less than 2kb) and are homologous at some level to the chromosomes, and, as such, represent paralogous/repeat sequences. For this reason, the unassigned KE supercontigs were not included in the initial CNV analysis. The full length *T. gondii* mtDNA genome is unassembled, and thus mtDNA sequence was not included in this first analysis.

A second *T. gondii* genome was curated that included mtDNA sequences as well as unique nonhomologous KE supercontigs. A portion of the *T. gondii* RH mtDNA genome spanning full length cytochrome b (*COB)* and cytochrome c oxidase subunit 1 (*COX1*) gene has been independently sequenced, GenBank: JX473253.1 (56). Although not complete, this 2607 bp sequence was added to the Tg14+api genome to represent the *T. gondii* mtDNA, named Tg14+api+mtDNA. The unassigned KE supercontigs were then compared to the Tg14+api+mtDNA genome using blastn and those KE supercontigs aligning with an expect value = 0.0 (80% of all KE supercontigs) were removed. The 455 remaining unique nonhomologous KE supercontigs were added to Tg14+api+mtDNA, resulting in a final curated *T. gondii* genome,Tg14+api+mtNDA+uniqueKE. This genome was used for bowtie2 alignment in a second analysis, revealing extensive CNV elevation across mtDNA sequence in the ART treated samples. SNP and indels were identified in mtDNA genes using VarScan (v2.4.3) (82) mpileup2snp or mpileup2indel commands with options --min-var-freq .05 and *P* value .05.

## Author contributions

AR, and LDS designed the research project; AR performed the experiments; AR, ML, MB, EW, and LDS analyzed data; AR and LDS wrote the paper with input from all authors.

## ACKNOWLEDGMENTS

We thank Jim Ajioka for advice on the mutation rate analyses, Sebastian Lourido for sharing unpublished data, Giel Van Dooren for providing TOM40 antibody, Jennifer Bark for technical assistance, and Wandy Beatty, Microbiology Imaging Facility, for assistance with electron microscopy. We thank the Genome Technology Access Center in the Department of Genetics at Washington University School of Medicine for assistance with genomic sequencing. The Center is partially supported by NCI Cancer Center Support Grant #P30 CA91842 to the Siteman Cancer Center and by ICTS/CTSA Grant UL1TR002345 from the National Center for Research Resources (NCRR), a component of the National Institutes of Health (NIH), and NIH Roadmap for Medical Research. This work was also supported by a grant from the NIH (AI118426) to LDS. AR was partially supported by a Berg Postdoctoral Fellowship from the Department of Molecular Microbiology at Washington University. ML was supported in part by a Ruth L. Kirschstein Institutional National Research Award from the National Institute for General Medical Sciences, T32 GM008666. This publication is solely the responsibility of the authors and does not necessarily represent the official view of NCRR or NIH.

