## Supplementary material for "Evolution of resistance in vitro reveals a novel mechanism of artemisinin activity in *Toxoplasma gondii*": Supplmental materials

**This PDF file includes:**

Supplementary text
Fig. S1
Tables S1 to S6
Additional Datasets S1 to S5

**Additional Dataset S1 (separate file)**
Mutational analysis for clones derived from the 12B line
**Additional Dataset S2 (separate file)**
Mutational analysis of clones derived from the unselected 5D line.
**Additional Dataset S3 (separate file)**
Mutational analysis of clones derived from the 100 uM ART selection from the B2 line
**Additional Dataset S4 (separate file)**
Mutational analysis of clones derived from the 100 uM ART selection from the F4 line
**Additional Dataset S5 (separate file)**
CNVs

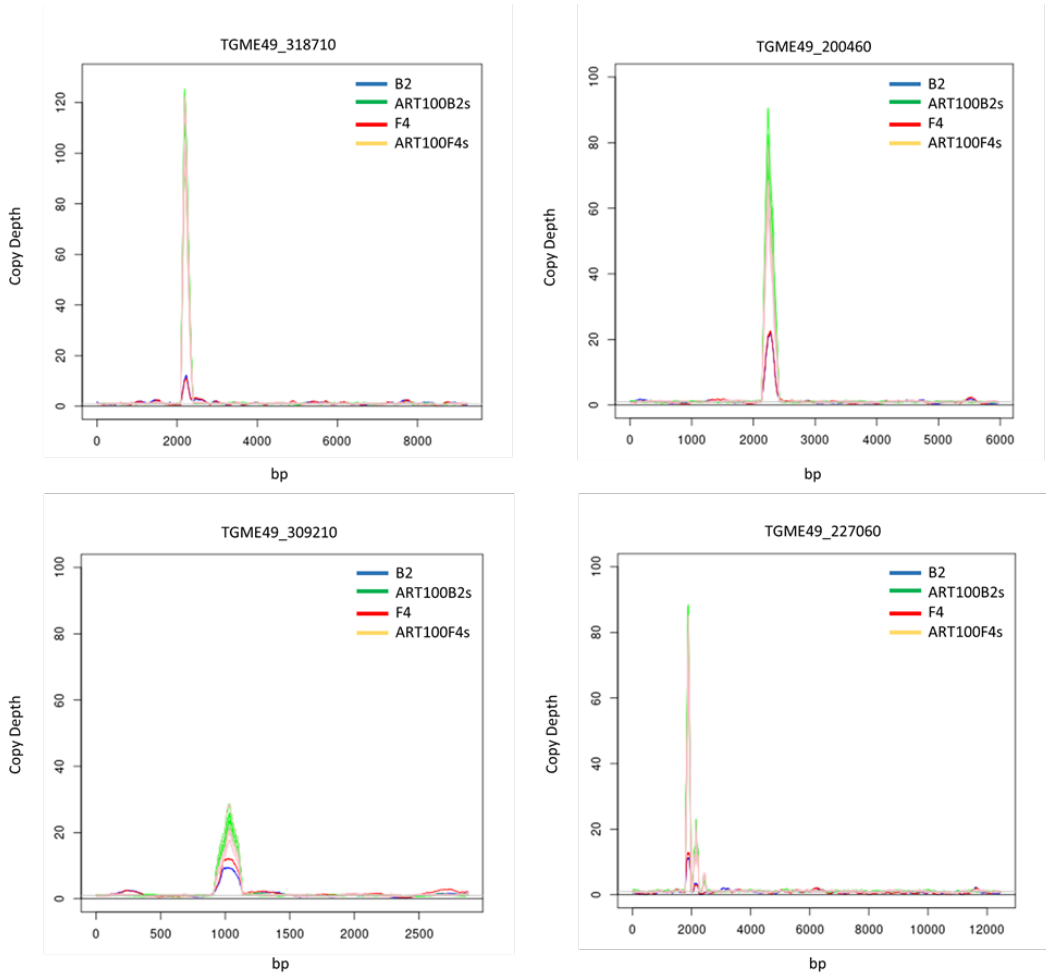


**Fig S1. Copy number variation depth for *T. gondii* strains across the detected CNV loci.** Copy number variation for wild type B2 parental, F4 parental and ART100-resistant lines derived from these parents. Illumina reads were mapped across *T.gondii* ME49 genome. The B2 line was sampled from the untreated (blue) or after extended passage at 100 µM (green) artemisinin. The F4 lines was sampled either untreated (red) or after extended passage at 100 µM (orange) artemisinin. Copy number estimates are based on the read depth per base pair normalized to 1X across the respective genome.

**Table S1 Mutations in candidate genes identified by whole-genome sequencing that were found only in one of the two ART resistant lines.**

| Line | Gene id[a] | Annotation | Frequency % (ART 8µM) | Frequency % (ART 16µM) | Frequency % (ART 100µM) | Coding region change | Amino acid change |
|---|---|---|---|---|---|---|---|
| B2 | TGME49_214110 | protein yipf5 | 0 | 97.78 | 100 | 578A>C | Glu193Ala |
| B2 | TGME49_310700 | serine/threonine phosphatase PP1 | 0 | 0 | 100 | 43C>G | Leu15Val |
| F4 | TGME49_263580 | bromodomain-containing protein | 0 | 0 | 100 | 1291C>A | Gln431Lys |
| F4 | TGME49_286030 | hypothetical protein | 6.25 | 100 | 100 | 1018T>G | Ser340Ala |
| F4 | TGME49_264030 | aminotransferase | 0 | 0 | 94.29 | 1553G>C | Arg518Pro |
| F4 | TGME49_202900 | zinc finger (CCCH type) motif-containing protein | 0 | 0 | 100 | 2083T>C | Trp695Arg |
| F4 | TGME49_314482 | hypothetical protein | 0 | 0 | 100 | 305C>G | Ala102Gly |

[a] www.ToxoDB.org

**Table S2. Growth Effects of ART on DegP and Ark1 point mutant strains**

| Strain | $EC_{50}$ µM |
|---|---|
| RHluc wild type | 0.220 ±0.019 |
| B2-DegP G806E | 0.267 ±0.023 |
| B2-Ark1 C274F | 0.272 ±0.021 |
| B2-DegP G806E/ Ark1 C274F | 0.307 ±0.03 |
| F4- DegP E821Q | 0.310 ±0.025 |
| F4- Ark1 C274R | 0.349 ±0.029 |
| F4- DegP E821Q/ Ark1 C274R | 0.356 ±0.028 |

The $EC_{50}$ data are presented as the average of three biological replicates (i.e., separate experiments) each with three internal replicates (i.e. separate wells).

**Table S3. CNVs detected in the ART 100µM selected lines.**

| Pools | Clones | Gene id[a] | Annotation |
|---|---|---|---|
| + | + | TGME49_200460 | Hypothetical protein |
| + | + | TGME49_318710 | ATP-binding cassette sub-family member 1 |
| + | partial | TGME49_309210 | Peroxiredoxin 6, putative |
| partial | partial | TGME49_227060 | Hypothetical protein |

[a] www.ToxoDB.org

**Table S4 Strains used in this study**

| Line | Genotype | Source |
|---|---|---|
| RH | RH | ATCC #50838 |
| RH-5D | RH | This study |
| RH-12B | RH | This study |
| RH-F4 | RH | This study |
| RH-B2 | RH | This study |
| RH*ΔΔ* | RH*Δhxgprt*Δ*ku80* | This study |
| RH*luc* | RH*ΔhxgprtΔku80*; *uprt*::*dhfr-ts*$^{[S36R, T83N]}$,TUB1:Firefly Luciferase | This study |
| RH DegP$^{[G806E]}$ | RH*ΔhxgprtΔku80*; *uprt*::*dhfr-ts*$^{[S36R, T83N]}$,TUB1:Firefly Luciferase; *degP*$^{[G806E]}$ | This study |
| RH DegP$^{[G821Q]}$ | RH*ΔhxgprtΔku80*; *uprt*::*dhfr-ts*$^{[S36R, T83N]}$,TUB1:Firefly Luciferase; *degP*$^{[G821Q]}$ | This study |
| RH Ark1$^{[C274F]}$ | RH*ΔhxgprtΔku80*; *uprt*::*dhfr-ts*$^{[S36R, T83N]}$,TUB1:Firefly Luciferase; *ark1*$^{[C274F]}$ | This study |
| RH Ark1$^{[C274R]}$ | RH*ΔhxgprtΔku80*; *uprt*::*dhfr-ts*$^{[S36R, T83N]}$,TUB1:Firefly Luciferase; *ark1*$^{[C274R]}$ | This study |
| RH DegP$^{[G806E]}$, Ark1$^{[C274R]}$ | RH*ΔhxgprtΔku80*; *uprt*::*dhfr-ts*$^{[S36R, T83N]}$,TUB1:Firefly Luciferase; *degP*$^{[G806E]}$, *ark1*$^{[C274R]}$ | This study |
| RH DegP$^{[G821Q]}$, Ark1$^{[C274F]}$ | RH*ΔhxgprtΔku80*; *uprt*::*dhfr-ts*$^{[S36R, T83N]}$,TUB1:Firefly Luciferase; *degP*$^{[G821Q]}$, *ark1*$^{[C274F]}$ | This study |

**Table S5 primers used in this study**

| Primer | Seq 5->3 |
|---|---|
| DegP G806E check/ amplify-F | GAAGGAGGTGGTTGACGTCC |
| DegP G806E check/ amplify-R | CTCCACTCGAGAGTTCGTGG |
| DegP G821Q check/ amplify-F | GATGGCTCGTGTCTTCTCTGCG |
| DegP G821Q check/ amplify-R | TATGGGTGGAGCCTGGCTTTG |
| Ark1 check/ amplify-F | GAGCATGCTCTTGTTGTGACC |
| Ark1 check/ amplify-R | TTGTTGGGTATGCCTGGCTC |
| Ark1 guide-F | GTCTGTTGACAGAGAAAACTGGTTTTAGAGCTAGAAATAGC |
| DegP G806E guide-F | GCTTGCAGGTTCCGCTTATCGGTTTTAGAGCTAGAAATAGC |
| DegP G821Q guide-F | GTTCTTCCTTCGCAAAGATGTGTTTTAGAGCTAGAAATAGC |
| Guide universal-L | AACTTGACATCCCCATTTAC |

**Table S6 gBlock sequences used as a template for Cas9-shielded homology donor amplicon to create the point mutant strains**

DegP G821Q gBlock®
GATGGCTCGTGTCTTCTCTGCGGATTTCGAGGCAAAGACTCCTTCAGTCTCTGACTGCCGGCTCCGCGTTTCGTCGCGGCCTCCGCGCTTT
CTGTGATCAACGGCTCGCGGGATTTCCGCGAGGAAACTCTGAGACCCGGGAGACACCCACGAACTCTCGAGTTGAGGACTAGTGGCCACC
GAGAGCTGGGGAAAACGCAGTGCCGGTTCACTCCCGTCTTCCTCGCAGGGACAGCAAACATGTTTTGTGTCTCTTCGTTCTTTCTCTCCGC
TTTTTCAGGATcAATTTGGAACGAAATTCAGCGAACGCGCACCTGCCTCCCTTCTCCAGCCTCTCGCaGAtATCTTTGCGAAGGAAGAAGGA
GAGGAACCTGTTATTCTCAGCCACGTAAGTCTCCTCATCTCAACACGTAAAACGACTCTCGAAACAAAGCCAGGCTCCACCCATA

Red lower case – ART resistance mutation
Highlighted sequence- guide RNA
Green lower case – silent mutations eliminating the PAM site and introducing an EcoRV restriction site.

DegP G806E gBlock®
GAAGGAGGTGGTTGACGTCCTTGTGAGTTCCACTTGTCACCCATTTCTTCAAGGAACAATGTCTTTCTTCCTCTTCTTCGGTGTCGCATCTTC
TCTCTTCTGTCTCCTTCTCCTTTCGTTTCTTTCCTGTCTTTTCGGTTCTCTCTTCTCTCTGTCTCTTTCTCTTTCCTTCTCTCTCATTTTTTCTCG
CCTGTTTCTCTTCTTCTCGCTTCTCTCTCCCCGTTTCCCTCTTCTGCGTTGGGTCTGTGTTGCCGGCACTCTTCGTTTTGCGGATATCTTTGC
TTGCAGGTTCCGCTTATCGAaGAGAACGCGCTTGTTCCGAAGCACCAGTGGGACAAGAAGGCGCGGTACCTCATCTACGaAGGACTTGTCT
TCTGTCCTCTCACTCTCGAaTACTTGAAGGTACGACCGAGTGGGCCGCAGATCTCAGATGGCCTCCCCCGCGATGGAGCGGCGCGTTGGA
CGGTTCCTCTCTCCCCTGTGGTTCTGATGGCTCGTGTCTTCTCTGCGGATTTCGAGGCAAAGACTCCTTCAGTCTCTGACTGCCGGCTCCG
CGTTTCGTCGCGGCCTCCGCGCTTTCTGTGATCAACGGCTCGCGGGATTTCCGCGAGGAAACTCTGAGACCCGGGAGACACCCACGAACT
CTCGAGTGGAG
Red lower case – ART resistance mutation
Highlighted sequence- guide RNA
Green lower case – silent mutation eliminating the PAM site.
Blue lower case – silent mutation eliminating ScaI restriction site.

Ark1 C274R gBlock®

GAGCATGCTCTTGTTGTGACCCACTCGAGATAAACGGGGTGTCGAAGTGGCAGCATCCGTATGCATTTTCATGTGTGTGCGGACCTACGCG
TCGCTTACGGCGTTTTTTGAATCGTGACGCTCCGTTTCTTCAGGCTTTGGCGACGCTTGTGCATGCGTTTGTTGAATCTTCCCGCTCACCGG
AATGTTTTGTCTCTGGACGCGATCTACGAGGAGAAACAGAAGTTCTACTTTGTTAGCGAGAAGCTCGAGGGCGGAGAGCTGTTTGATTTTCT
GTTGACAGAGAAAACTGTcGAGGAGCACATAcGCCAGTACATCATCTTCCAGATTTTGCAGGCTCTCAATCACATGCACTCGAACCACCTTC
TTCACAGGTGAGAAGACGCAGAATCGCGGGCGTCTCGCCTTGCATCTGACGGTCAAATATCGCTTTCAACATGATCATGTTCAAGAACAAG
GAACTAATGAGCCGACATCGATATGTAGATACAGGCCGAGGGATACAGGTCCCCACAGTAACTGACCTGTTTTACCAGGCGTGAGATACGC
CCGGAAAACAAGTTAGAAACAGAATGCTTTAAGACAAACTGCGGTGTATGGGAGCCAGGCATACCCAACAA

Red lower case – ART resistance mutation and eliminating NdeI restriction site.

Highlighted sequence- guide RNA

Green lower case – silent mutations eliminating the PAM site.

Ark1 C274F gBlock®

GAGCATGCTCTTGTTGTGACCCACTCGAGATAAACGGGGTGTCGAAGTGGCAGCATCCGTATGCATTTTCATGTGTGTGCGGACCTACGCG
TCGCTTACGGCGTTTTTTGAATCGTGACGCTCCGTTTCTTCAGGCTTTGGCGACGCTTGTGCATGCGTTTGTTGAATCTTCCCGCTCACCGG
AATGTTTTGTCTCTGGACGCGATCTACGAGGAGAAACAGAAGTTCTACTTTGTTAGCGAGAAGCTCGAGGGCGGAGAGCTGTTTGATTTTCT
GTTGACAGAGAAAACTGTcGAGGAGCACATATgCCAGTACATCATCTTCCAGATTTTGCAGGCTCTCAATCACATGCACTCGAACCACCTTC
TTCACAGGTGAGAAGACGCAGAATCGCGGGCGTCTCGCCTTGCATCTGACGGTCAAATATCGCTTTCAACATGATCATGTTCAAGAACAAG
GAACTAATGAGCCGACATCGATATGTAGATACAGGCCGAGGGATACAGGTCCCCACAGTAACTGACCTGTTTTACCAGGCGTGAGATACGC
CCGGAAAACAAGTTAGAAACAGAATGCTTTAAGACAAACTGCGGTGTATGGGAGCCAGGCATACCCAACAA

Red lower case – ART resistance mutation and eliminating NdeI restriction site.

Highlighted sequence- guide RNA

Green lower case – silent mutations eliminating the PAM site.
